# Alteration of Vascular Endothelial Cadherin in Alzheimer’s Disease Patient and Mouse Model

**DOI:** 10.1101/430140

**Authors:** Daehoon Lee, Sun-Jung Cho, Hyun Joung Lim, JiWoong Seok, Chulman Jo, Sangmee A. Jo, Moon Ho Park, Changsu Han, Neil Kowall, Hoon Ryu, Rudolph E. Tanzi, Young Ho Koh

## Abstract

Alzheimer’s disease (AD) is characterized by amyloid plaques and pathologic cerebrovascular remodeling. Cerebrovascular abnormalities may contribute to the pathology of AD, but the molecular mechanisms are not fully understood. In this study, we evaluated blood–brain barrier (BBB) disruption and the role of VE-cadherin in the progression of amyloid pathology. Here, we determined that levels of VE-cadherin are decreased in brain vessels of AD patients and mouse model of AD. *In vitro* experiments showed that the disappearance of VE-cadherin by beta-amyloid at the endothelial cell surface was due to cleavage of VE-cadherin. VE-cadherin cleavage was inhibited by a γ-secretase and ADAM10 inhibitor. The disappearance of VE-cadherin in brain vessels was also seen in amyloid precursor protein transgenic mice. In the postmortem brain of individuals with AD, furthermore, levels of VE-cadherin were significantly reduced in vessels. Dementia patients showed a distinct blood biochemical profile characterized by high soluble VE-cadherin (sVEC). There was a strong association between plasma sVEC (adjusted odds ratio = 3.41, *P* < 0.001) and dementia. These results suggest that measurements of plasma VE-cadherin could have the potential for predicting the risk of progressive AD.

## Introduction

Alzheimer’s disease (AD) is the brain disease marked by progressive cognitive decline and is characterized by the accumulation of beta-amyloid (Aβ) deposits and neurofibril tangles (Joachim et al, 1989; Tanzi et al, 1992). The major component of senile plaques and cerebrovascular deposition is the 39–43 residue polypeptide derived by proteolytic processing of amyloid precursor protein (APP) (Gillespie et al, 1992). Aβ plays a critical role in the development of AD pathology (Deane & Zlokovic, 2007; Selkoe, 2001; Tanzi, 2005). Several studies have found considerable abnormalities and deficiencies in properties of the blood-brain barrier (BBB) in the AD brain (Zipser et al, 2007). Lesions in the white matter have been observed in the AD brain(Brun & Englund, 1986). Therefore, microvascular lesions have also been postulated to be a common pathological feature appearing during the development of AD (Brun & Englund, 1986; White et al, 2002). Amyloid accumulation may be closely associated with cerebrovascular damage.

The BBB is a tightly controlled semi-permeable barrier that protects the brain from toxic substances in the circulating blood (Poduslo et al, 1994). The BBB is formed by pericytes, astrocytes and capillary endothelial cells (ECs) packed by tight junctions (TJs) and adherens junctions (AJs). These cells play a significant role in BBB integrity, and disruption of the cadherin junctions between pericytes and endothelial cells has recently been shown to disrupt the BBB (Li et al, 2011). Increased BBB dysfunction has been reported CNS diseases including AD (Erickson & Banks, 2013). Although BBB impairment might play an important role in the pathogenesis of AD, it remains unclear whether a BBB disruption is a part of the pathophysiology of AD. Recent post-mortem studies have shown that BBB is disrupted in AD brain tissue as compared to controls that is associated with degeneration of pericytes (Sengillo et al, 2013). Recent neurovascular studies also revealed that Aβ contributes to injury resulting in BBB leakage, including cortical microbleeds in humans (Hartz et al, 2012). In addition, Aβ also contributes to decreased levels of TJ components, such as ZO-1 (Kook et al, 2012; Marco & Skaper, 2006).

At the endothelial AJs in the brain of vertebrates, adhesion is mediated by the integral membrane protein vascular endothelial cadherin (VE-cadherin; VEC also known as CD144) (Vestweber et al, 2009). VEC consists of one extracellular fragment with five domains, one short transmembrane domain, and a cytoplasmic domain that interacts with catenins (Lampugnani et al, 1995). VEC is important to control vascular permeability both *in vitro* and *in vivo*(Dejana et al, 2008). Blocking antibodies to VEC induce a marked increase in human umbilical vein endothelial cells (HUVECs) and lung permeability accompanied by hemorrhage (Corada et al, 2001). Recently, it was reported that the ectodomain of VEC could be released from endothelial cells, suggesting that VEC cleavage may play a role in reorganization of AJs (Herren et al, 1998; Schulz et al, 2008). Vascular endothelial growth factor receptor-2 (VEGFR2) can associate with VE-cadherin to facilitate the phosphorylation of AJ components by Src, and might thereby induce endothelial dysfunction (Dejana et al, 2008). Recently we have reported that the levels of VEGFR2 were decreased in the brains of AD model mouse and Aβ-treated endothelial cells (Cho et al, 2017). While the role of tight junctions is important to understand BBB hyperpermeability, few studies have addressed the role of AJs in amyloid-mediated endothelial dysfunction in AD. Although impairment of this barrier by Aβ deposits leads to disintegration of endothelial junctional and an increase in permeability of the BBB, little is known about the molecular mechanisms regulating the cohesion of endothelial AJs in AD. Moreover, these structural modifications of VEC have not been explored.

Here, we identified a role of Aβ-induced VEC adhesion junction disruption in AD. We propose that an initial increase in cerebral Aβ deposits leads to BBB disruption. This triggers a vicious cycle of BBB damage, which has a causal relationship in the AD mouse brain. Our data provide evidence for alteration of VEC in the brain of AD patients. In addition, soluble VE-cadherin (sVEC) levels in blood were elevated in dementia patients. Our results suggest that alteration of VE-cadherin may reflect the attenuation of BBB integrity in AD. Plasma VE-cadherin may have potential as a useful blood-based biomarker of microvascular pathology in AD.

## Results

### Expression of VEC is decreased at the cerebral capillaries in the cortex of Alzheimer’s brain

Congophilic amyloid plaques first appear at 5 months and increase in number with age in APP Swedish/PS1dE9 transgenic (APP Tg) mice (Fig EV1A). To assess the changes in cerebral capillaries in the APP Tg mouse, we performed double staining with anti-collagen type IV (Coll-IV) and anti-Aβ, 6E10. Coll-IV was used as a marker of basal lamina damage and BBB disruption (Trinkl et al, 2006). Cerebral blood vessels stained for Coll-IV showed long tubular-like form, and 6E10 immunoreactivity was not detected in the brain of wild-type mice. Severe destruction of capillaries was observed near the areas of amyloid plaque deposition in the cortex of 21-month-old APP Tg mice (Fig EV2A, white dotted circle), while there was no capillary disruption near such areas in the cortex of 9-month-old APP Tg mice (Fig EV2B). To monitor BBB breakdown, we examined the presence of IgG in the brain parenchyma. We found that IgG leakages in brain were detected in APP Tg mice (Fig EV3A).

We next performed double staining with anti-VE-cadherin (BV9) and anti-Coll-IV antibodies. VEC was expressed in cerebral capillaries in the wild-type (Fig 1A). In contrast, VEC expression was markedly reduced in the brain capillaries of 21-month-old APPTg mice, even though Coll-IV expression partially remained at the basal lamina of the BBB (Fig 1A). As there was no capillary destruction near the areas of amyloid plaque deposition in the 9-month-old APP Tg mice (Fig EV2B), we also investigated the disappearance of VEC in the brains of 2-, 9-, and 12-month-old APP Tg mice (Fig EV4). Occasional VEC disappearance was seen by 9 months old with abundant plaques and a progressive decrease in VEC expression by up to 12 months. In order to assess the clinical significance of VEC alteration in vessels, we obtained entorhinal cortex region of postmortem brain samples from AD patients and control human subjects. The disappearance of VEC in vessel of brain tissue was also observed in AD patients (Fig 1B).

**Figure 1.**
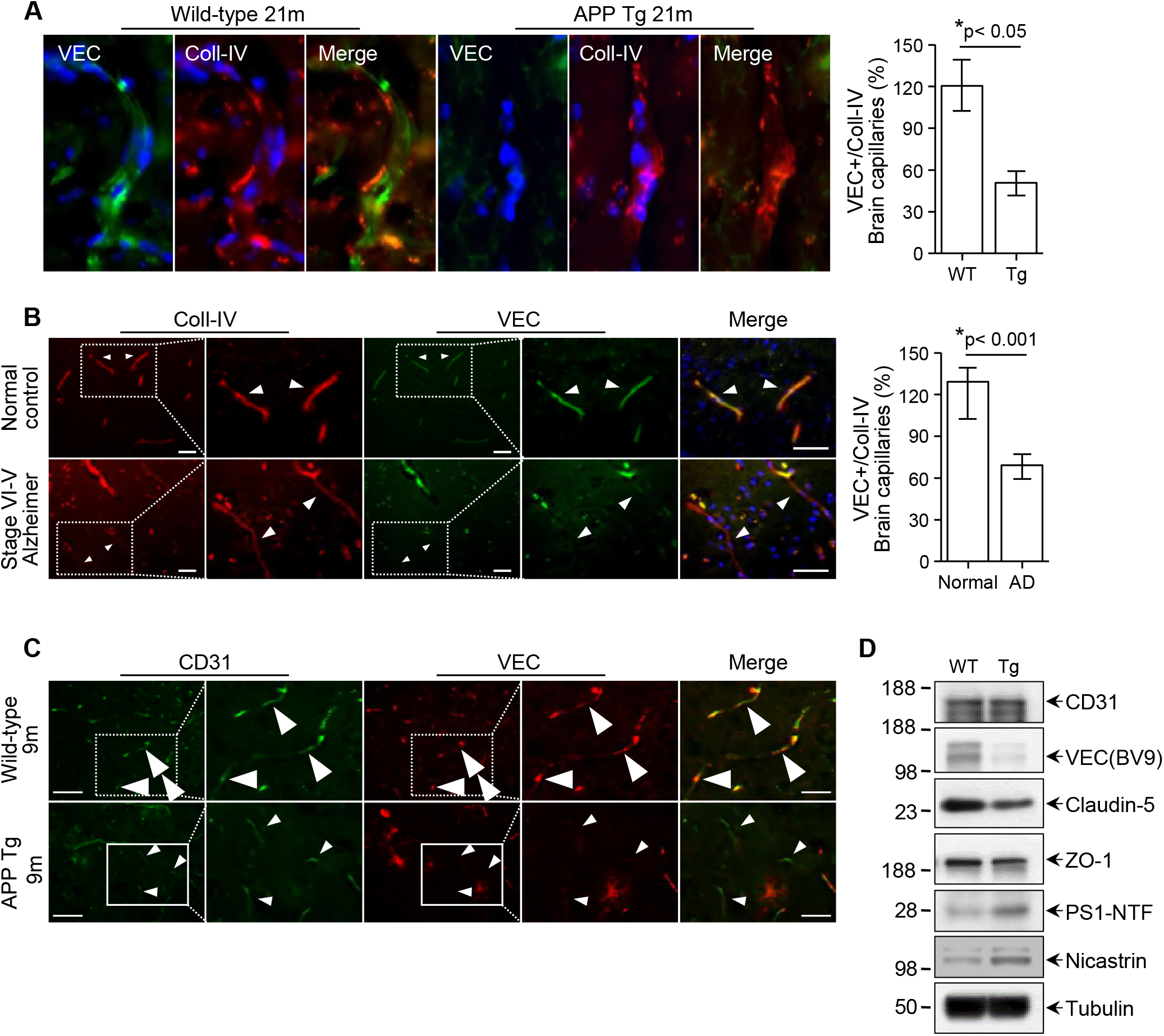
VE-cadherin (VEC) expression is decreased in the Alzheimer’s Brain. **A**. Immunostainings of VEC (BV9) and Coll-IV in the brain of 21-month-old wild-type and APP Tg, High-magnification images of cerebral capillaries, DAPI staining for nucleus, scale bar, 50 μm. Quantification of VEC-positive signal expressed as VEC-positive area (%) occupying Coll-IV-positive endothelial capillary profiles in the wild type and APP Tg mice. Mean ± SEM, n = 3 mice per group, \**P* < 0.05. **B**. Immunohistochemistry of VEC (BV9) and Coll-IV in the entorhinal cortex of human brain, Lower-magnification images scale bar, 50 μm, and high-magnification images scale bar 100 μm. Quantification of VEC-positive immunofluorescent signal normalized by Coll-IV-positive capillaries in the entorhinal cortex of human brain. Mean ± SEM, n = 3 postmortem specimen per group, * *P* < 0.001. **C**. Immunostainings of both CD31 and VEC (BV9) in the cerebral capillaries of 9-month-old wild-type and APP Tg mice brain, scale bar, 50 μm. High-magnification images of cerebral capillaries, scale bar, 100 μm. Arrowheads indicate alterations in VEC distribution in APP TG and human specimen. **D**. Immunoblots were performed with anti-CD31, VEC (BV9), claudin-5, ZO-1, presenilin1 N-terminal fragment (PS1-NTF) and nicastrin in the cerebral cortex of 10-month-old wild-type and APP Tg mice (n = 3). Anti-Tubulin was used as a loading control.

To monitor alterations of AJs in the brains of APP Tg mice, we also investigated expression of the endothelial marker platelet endothelial cell adhesion molecule (PECAM-1, CD31) in the brain (Fig 1C and Fig EV5). Reduced CD31 expression was observed in some areas of the brains of 9-month-old APP Tg mice and became more extensive with age (21-months old) (Fig EV5). The level of CD31 decreased where amyloid plaques abutted the perivascular space in the microvessels. Interestingly, reduced VEC immunoreactivity was observed in some cerebral vessels expressing CD31 (Fig 1C). We next investigated alterations of TJ-associated protein in APP Tg mice (Fig EV6). In some cerebral vessels in APP Tg mice, immunohistochemical staining of the TJ proteins including occludin (Fig EV6A) and ZO-1 (Fig EV6B) was not changed, but that of VEC protein was significantly reduced. These observations suggest that VEC may be more vulnerable and easily reduced on cerebral vessels of APP Tg mice. We next examined whether the levels of BBB-associated proteins are altered in APP Tg mice. Western blotting analysis was performed using brain lysates of 10-month-old wild-type and APP Tg mice. As expected, the levels of TJ-related proteins, such as ZO-1, were decreased in the APP Tg mouse brain compared with wild-type controls. We found a significant decrease in VEC protein level in the APP Tg mouse brain compared with wild-type controls (Fig 1D).

### Aβ_1–42_ alters disruption of VEC in human endothelial cells

To investigate the molecular mechanism underlying the reduction of VEC levels on vessels of APP Tg mice, HUVECs were treated with various concentration of Aβ_1–42_ for 24 and 48 h. As expected, the levels of full-length VE-cadherin (VEC/FL) were reduced after treatment with Aβ_1–42_ (Fig 2 A-B and Fig EV7A). Aβ treatment of HUVECs resulted in time-dependent (Fig 2A) and dose-dependent (Fig EV7A) productions of VEC cleavage product (VEC/CTF1). However, treatment with Aβ_42–1_ peptides did not affect VEC cleavage (Fig EV7B). These results indicated that Aβ_1–42_ is a potent inducer of VEC cleavage. We next examined secretion of the N-terminal fragment of VEC (soluble VE-cadherin; sVEC) into the conditioned medium. sVEC has a molecular mass of about 90 kDa and was detected in untreated cells. The levels of sVEC in conditioned medium increased after Aβ_1–42_ treatment (Fig 2B and Fig EV7A). The Aβ-mediated cleavage of VEC cytoplasmic domains may lead to the release of VEC extracellular domain (sVEC) and would be expected to decrease the VEC level on the cell surface. To examine this possibility, we performed immunofluorescence microscopy. Staining for VEC at the plasma membrane of HUVECs with anti-VEC (BV9) was reduced after treatment with Aβ (Fig 2C).

**Figure 2.**
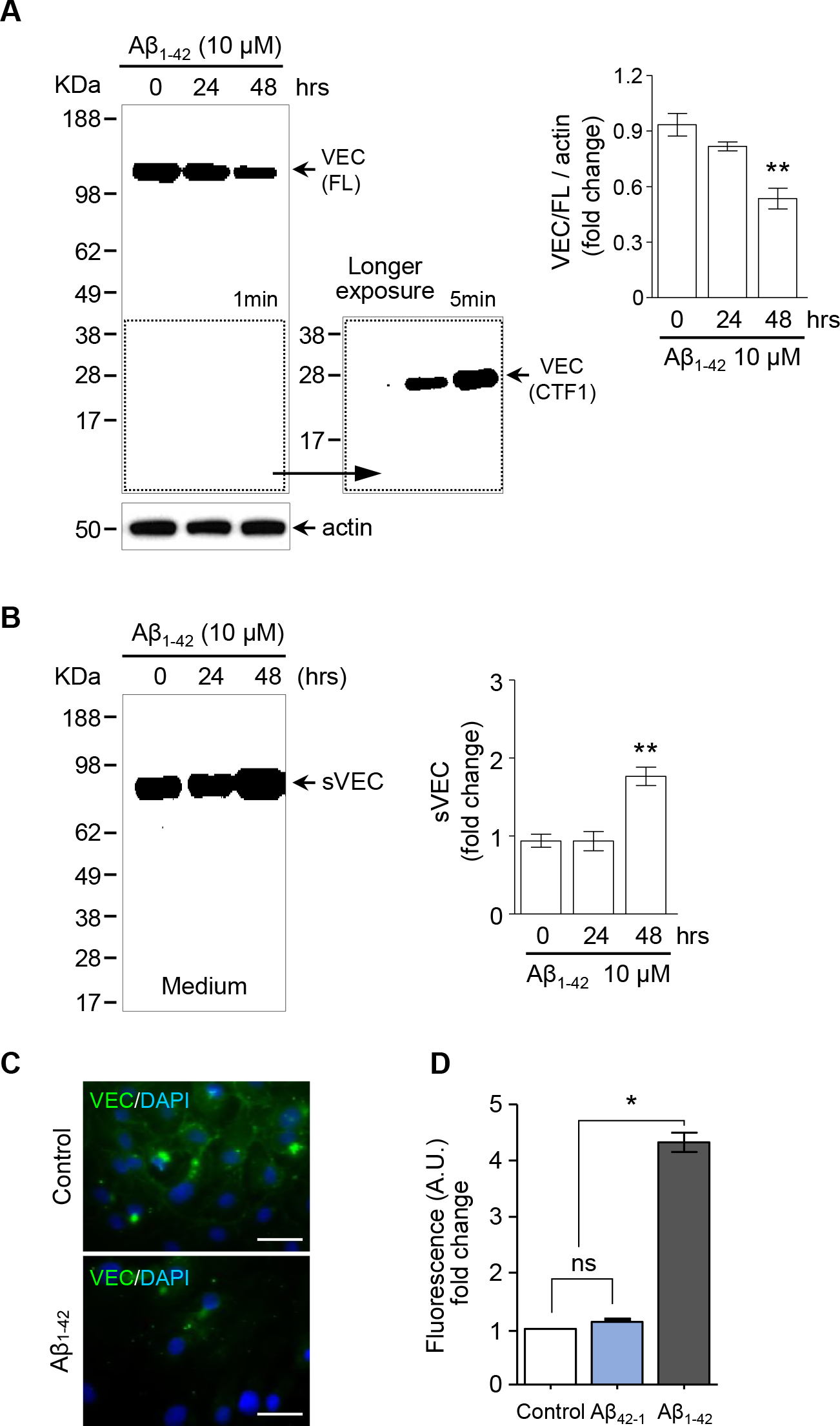
Aβ alters expression of VEC in human endothelial cells. **A**. HUVECs were treated for 0, 24, and 48 h with 10 μM Aβ_1–42_. Total lysates were immunoblotted with a rabbit anti–VE-cadherin (corresponding to the c-terminus). Quantification of VEC/FL fold change by densitometry, \*\**P* < 0.01 (student’s t-test). **B**. The medium was collected, and western blotting was performed using mouse anti-VEC (BV9; corresponding to the ectodomain). Quantification of sVEC fold change by densitometry, \*\**P* < 0.01 (student’s t-test). **C**. HUVECs were grown to confluence on collagen-coated glass coverslips and treated with Aβ_1–42_(10μM). Cells were immunostained with anti-VEC (BV9) and nuclei were counterstained with DAPI. Scale bar represents 20μm. After treatment of Aβ_1–42_ or nonamyloidogenic scrambled Aβ_42–1_ peptides in HUVECs, changes of diffusion of HUVECs monolayer were measured on 48 h by FD-40. Data are represented as mean ± SEM of three independent experiments performed in triplicate. \*\*\**P*< 0.001 versus control sample.

To evaluate whether the decrease in VEC level in Aβ-treated HUVECs reflected a change of endothelial cell-cell junctions, HUVECs were cultured in a Transwell system to measure released FITC-dextran 40 (FD-40). Treatment of 10 μM Aβ_1–42_ induced a significantly greater increase in FD-40 level in the plate chambers than control or Aβ_42–1_(Fig 2D). When HUVECs were treated with Aβ peptides, cell viability was not reduced by 15 μM of both Aβ_1–42_ and Aβ_42–1_(Fig EV7C and D). These results indicated that Aβ_1–42_ disrupts endothelial cell-cell junctions and increases endothelial permeability.

### Aβ induces the release of the VEC ectodomain via γ-secretase and ADAM 10

It was reported previously that γ-secretase was involved in the processing of cadherin (Marambaud et al, 2003). Therefore, we further investigated whether the cleavage of VEC by Aβ is followed by γ-secretase-mediated processing. The γ-secretase inhibitor DAPT treatment reduced the level of sVEC increased by Aβ, indicating a role of the γ-secretase in Aβ-mediated VEC cleavage (Fig 3 A). We measured the permeability of HUVECs to examine whether γsecretase affects the integrity of AJs. Culture on Transwell filter inserts together with DAPT resulted in a significant decrease of endothelial permeability compared to Aβ-treated cells (Fig 3B). Increased endothelial permeability is associated with impaired intercellular contact. Staining for VEC at the plasma membrane of HUVECs with anti-VEC (BV9) showed that pre-incubation with DAPT significantly delayed the loss of VEC at the cell surface after Aβ treatment (Fig 3C). Exposure of HUVECs to Aβ for 72 h decreased the levels of TJ proteins, such as ZO-1, occludin, and claudin-5 (Fig EV8A). Staining for VEC at the plasma membrane of HUVECs with anti-VEC antibody was reduced after treatment with Aβ for 48 h (Fig EV8B). Preincubation with DAPT significantly delayed the loss of VEC at the cell surface after Aβ exposure. However, treatment with Aβ for 48 h did not reduce expression of ZO-1 at the HUVEC plasma membrane (Fig EV8C). Moreover, reduced ZO-1 levels at the surface of HUVECs were detected after treatment with Aβ for 72 h. Preincubation with DAPT also delayed the loss of ZO-1 staining at the cell surface after Aβ exposure (Fig EV8D). In addition, we also examined a role of other proteases in VE-cadherin proteolysis. When EC was pretreated with ADAM10 inhibitor GI254023X, Aβ-mediated sVEC production was blocked by ADAM10 inhibitor GI254023X, not MMP inhibitor GM6001 (Fig EV9A). Furthermore, the levels of ADAM10 mRNA were significantly increased in HUVEC by Aβ treatment for 8 h (Fig EV9B). It is well known that cadherin cleavage was induced by calcium which is increased by Aβ in the cells (Kawahara et al, 2000; Park et al, 2008). We also found that Aβ-mediated sVEC production was blocked by treatment of 10 μM intracellular calcium chelator BAPTA-AM (Fig EV9C). VEGF activation of EC results in phosphorylation and disassembly of VEC (Dejana et al, 2001). As tyrosine phosphorylation of VEC-CTF is involved in increased permeability by VE-cadherin cleavage (Dejana et al, 2008; Vilgrain et al, 2013), we analyzed VE-cadherin tyrosine phosphorylation. The phosphorylation on Y^685^ VE-cadherin was decreased in HUVEC after Aβ treatment (Fig 3D).

**Figure 3.**
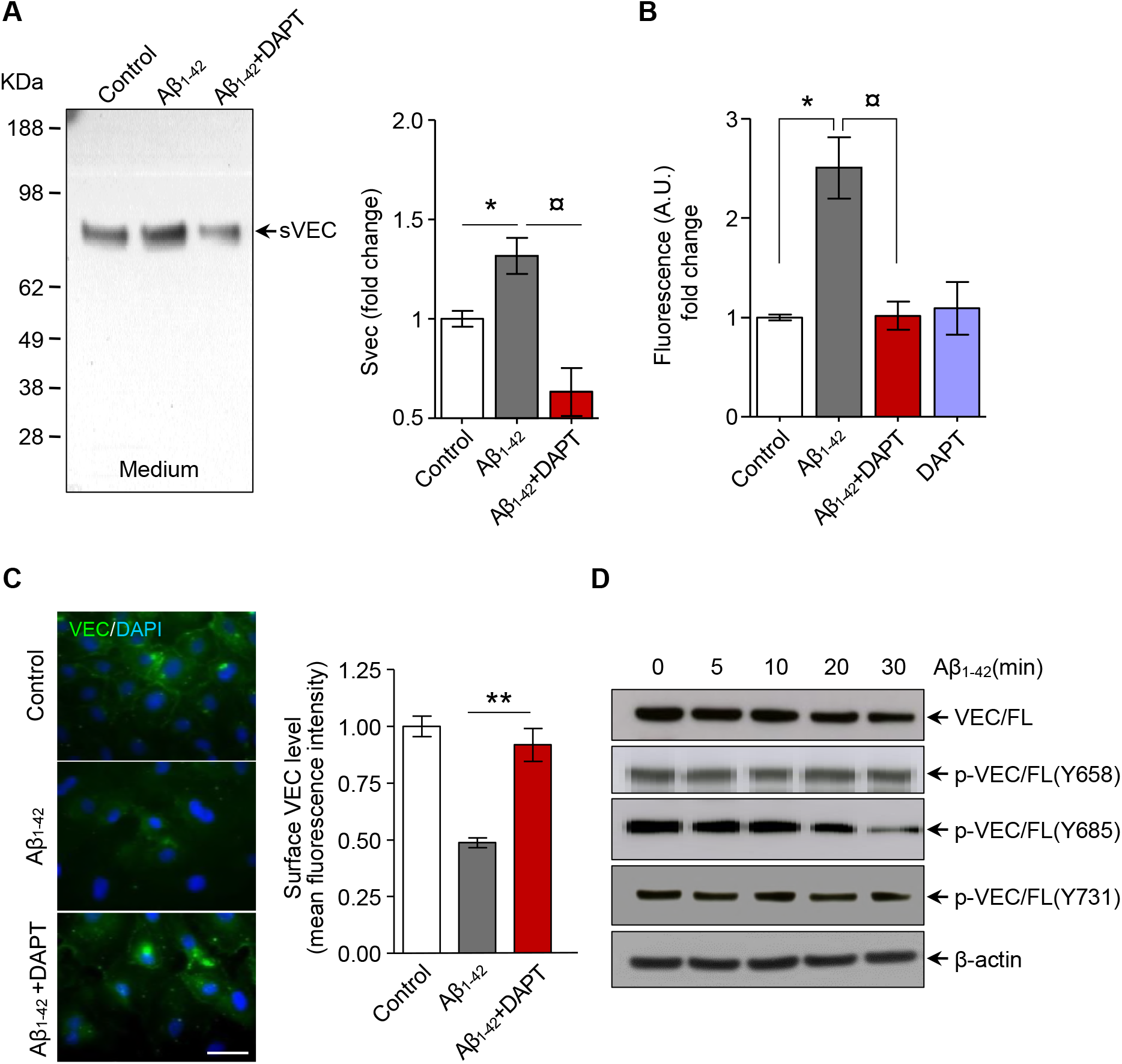
γ-secretase inhibitor prevents Aβ_1–42_-induced dissociation of endothelial VEC. **A**. After pretreatment with DMSO or γ-secretase inhibitor DAPT (2.5 μM) for 1 h, HUVEC cells were treated with Aβ_1–42_ (10 μM) for 48 h. The medium was collected, and western blotting was performed using anti-VEC (BV9). Quantification of sVEC folds changes by densitometry. **B**. After pretreatment with DMSO or γ-secretase inhibitor DAPT (2.5 μM) for 1 h, HUVEC cells were treated with Aβ_1–42_ (10 μM). Changes of diffusion of HUVECs measured in 48 hours by FD-40 were measured under the same condition as perform **Fig 2D**. Data are represented as mean ± SEM expressed as mean; n = 3 independent experiments. \**P*< 0.05, versus control sample; ¤*P*< 0.05 versus Aβ_1–42_-treated sample. **C**. Cells were immunostained with VEC (BV9) and nuclei were counterstained with DAPI. Scale bar represents 20 μm. Quantification of surface levels of VEC fold change by densitometry. Data are represented as mean ± SEM expressed as mean; n = 3 independent experiments. \*\**P* < 0.01, versus Aβ_1–42_-treated sample. **D**. AfterHUVEC cells were treated with Aβ_1–42_ (10 μM) for various times, total lysates were immunoblotted with a anti–VE-cadherin (BV9), anti-phospho VE-cadherin (Y658, Y685, Y731).

The phosphorylation on Y^658^ and Y^731^ VE-cadherin were not changed after Aβ treatment. Our results implicate that Aβ induces VE-cadherin proteolysis and increase sVEC levels via γsecretase and ADAM10.

### Soluble VEC is found in amyloid plaque

As the cleavage of VEC in brain vessels could lead to release of sVEC into the blood, we next examined whether sVEC is associated with amyloid plaques. We performed immunostaining of brain sections from wild-type and APP Tg mice. Unlike vWF staining (Fig EV10A), extravascular deposition of VEC was enhanced in the cortex of APP Tg mouse brain sections and colocalized with some of the senile plaques stained for thioflavin-S (Fig 4A and Fig EV11A). There was no detectable VEC immunoreactivity with the intracellular VEC antibody (770–781) (Fig EV10B). Furthermore, in the amyloid plaques stained with anti-Aβ_1–42_, there was a massive increase in reactivity for sVEC (Fig 4B). Our results demonstrate considerable extents of sVEC colocalized with amyloid plaques.

**Figure 4.**
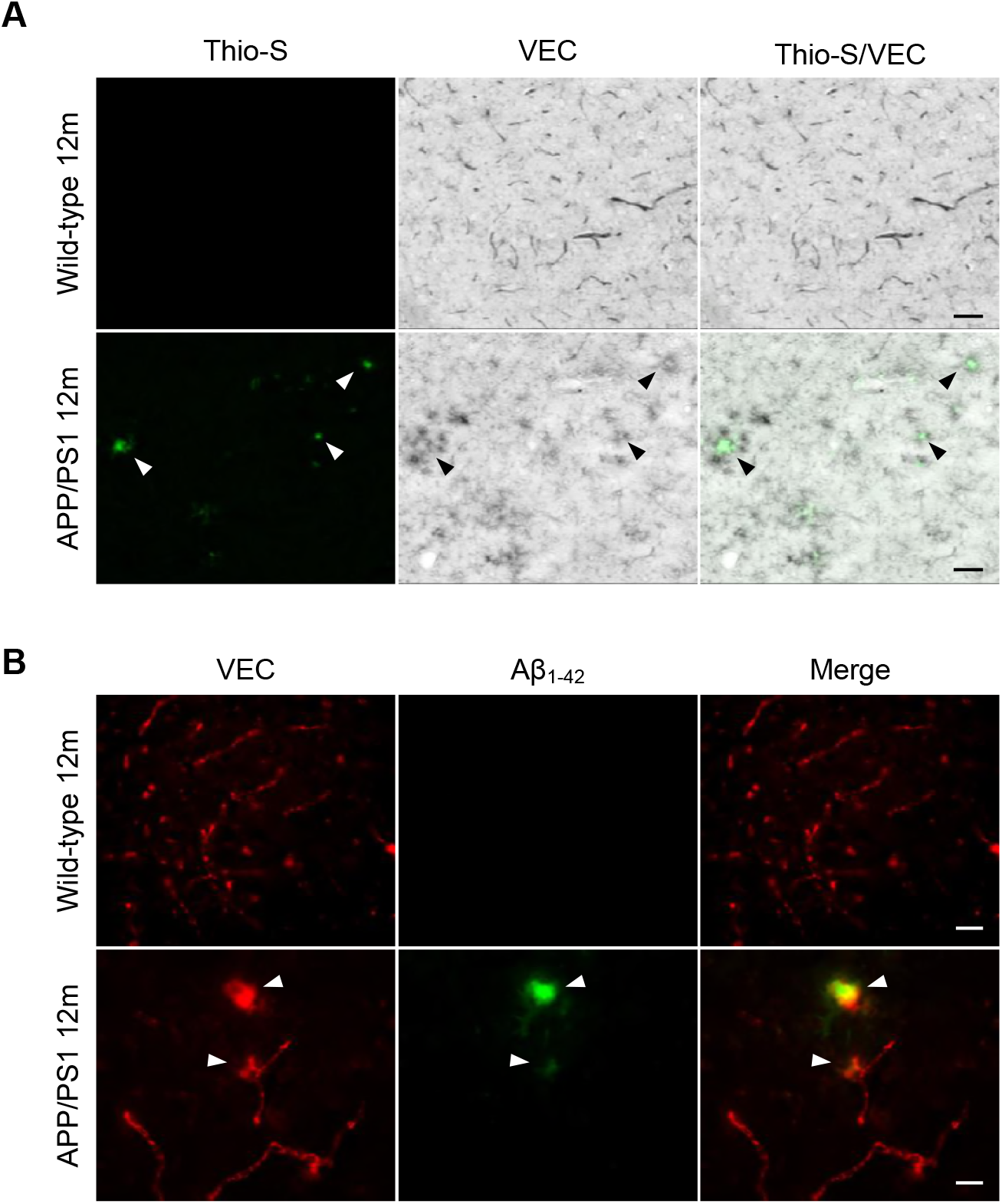
sVEC is colocalized with amyloid plaques in AD mouse brain. **A**. Sections from the cortex of 12-month-old wild-type and APP Tg mice were immunolabeled with anti-VEC (BV9) and counterstained with thioflavin-S for amyloid plaques (Scale bar represents 250 μm). **B**. Representative immunofluorescence of anti-VEC (BV9) with anti-Aβ_1–42_ in 12-month-old wild-type and APP Tg mice (Scale bar represents 20 μm). Arrowheads indicate that sVEC immunoreactivity was found in the amyloid plaques. Results are representative of three mice per group from two separate experiments.

### Plasma soluble VEC level is elevated in Alzheimer’s disease patients

For a more indication of endothelial dysfunction in AD pathogenesis, we measured sVEC in plasma from wild-type and APP Tg mice. As VEC disappeared at the cell surface in the brains of APP Tg mice, sVEC levels in the blood of these mice may be elevated. As expected, the blood sVEC levels were markedly elevated in 10-month-old APP Tg mice (Fig 5A). We next investigated the alterations in sVEC level with aging. We first collected plasma from 7-month-old APP Tg mice and wild-type controls. After 2 months, blood was collected again from the same mice. The sVEC levels in these two plasma samples from each wild-type and APP Tg mouse were measured by ELISA. In APP Tg mice, the plasma sVEC levels in 9-month-old mice were elevated compared to those at 7 months old (Fig 5B). We also examined the levels of adhesion molecules (VCAM-1 and ICAM-1) and metalloproteinase (ADAM10 and MMP9) in plasma from 18-month-old wild-type and APP Tg mice. There was no alteration of plasma levels of VCAM-1, ICAM-1, ADAM10 and MMP9 in mice (Fig EV12A). To further support our findings, we measured sVEC levels in the plasma of human dementia patients. To clarify whether levels of sVEC are related to clinically overt dementia, we examined sVEC levels in MCI and dementia patients as well as in controls without dementia (Fig EV13A). The clinical characteristics of MCI and dementia patients and controls are summarized in Table EV1. Western blotting analysis indicated that sVEC levels were increased in the plasma of dementia patients (n = 3) compared to controls without dementia (n = 3) (Fig EV13 B and C). For analysis of sVEC concentration in human plasma, we determined sVEC levels by ELISA. The study flow diagram in Fig EV13A provides further details on sample use. We first examined 150 plasma samples from set1 (validation study). We also examined clinical pathology variables, several biomarkers and 1 metal ion (Table EV3). The levels of three biomarkers (sVEC, VCAM-1, and cholesterol) were different between the three groups (*P*< 0.05; Kruskal-Wallis test). The levels of four biomarkers (sVEC, VCAM-1, HDL, and homocysteine) were different between control and dementia groups (*P*< 0.05; Mann-Whitney U test). The levels of three biomarkers (VCAM-1, cholesterol, homocysteine) were different between MCI and dementia groups (*P*< 0.05; Mann-Whitney U test). Interestingly, plasma sVEC levels only were different between control and MCI groups (*P*< 0.05; Mann-Whitney U test). Table EV4 indicates that there was a negative correlation between the plasma level of sVEC and the results of the MMSE assessments. However, the CDR assessment was positively correlated with the plasma level of sVEC. There was no correlation between the plasma level of sVEC and biomarkers of microvascular pathology (ICAM-1, VCAM-1), and the clinical biomarkers of cardiovascular risk (total cholesterol, triglycerides, etc.) (Table EV5). In the verification study (Set1 + Set2), plasma sVEC concentrations were also elevated in MCI subjects or Dementia patients compared to the control subjects (*P*< 0.001; Mann-Whitney U test) (Fig 5C and Table EV6). The receiver operating characteristic (ROC) curve were run to evaluate specificity and sensitivity of sVEC, to distinguish dementia from control subjects. The AUC of ROC curve on Dementia versus control using plasma sVEC was 0.72 (*P*< 0.05) (Fig 5D). Logistic regression was used to estimate the odds ratio (ORs) and 95% CIs for (Table EV6). Plasma sVEC was associated with more than a 3.4-fold increase in the risk of Dementia. Our data suggested that plasma sVEC could be elevated in AD.

**Figure 5.**
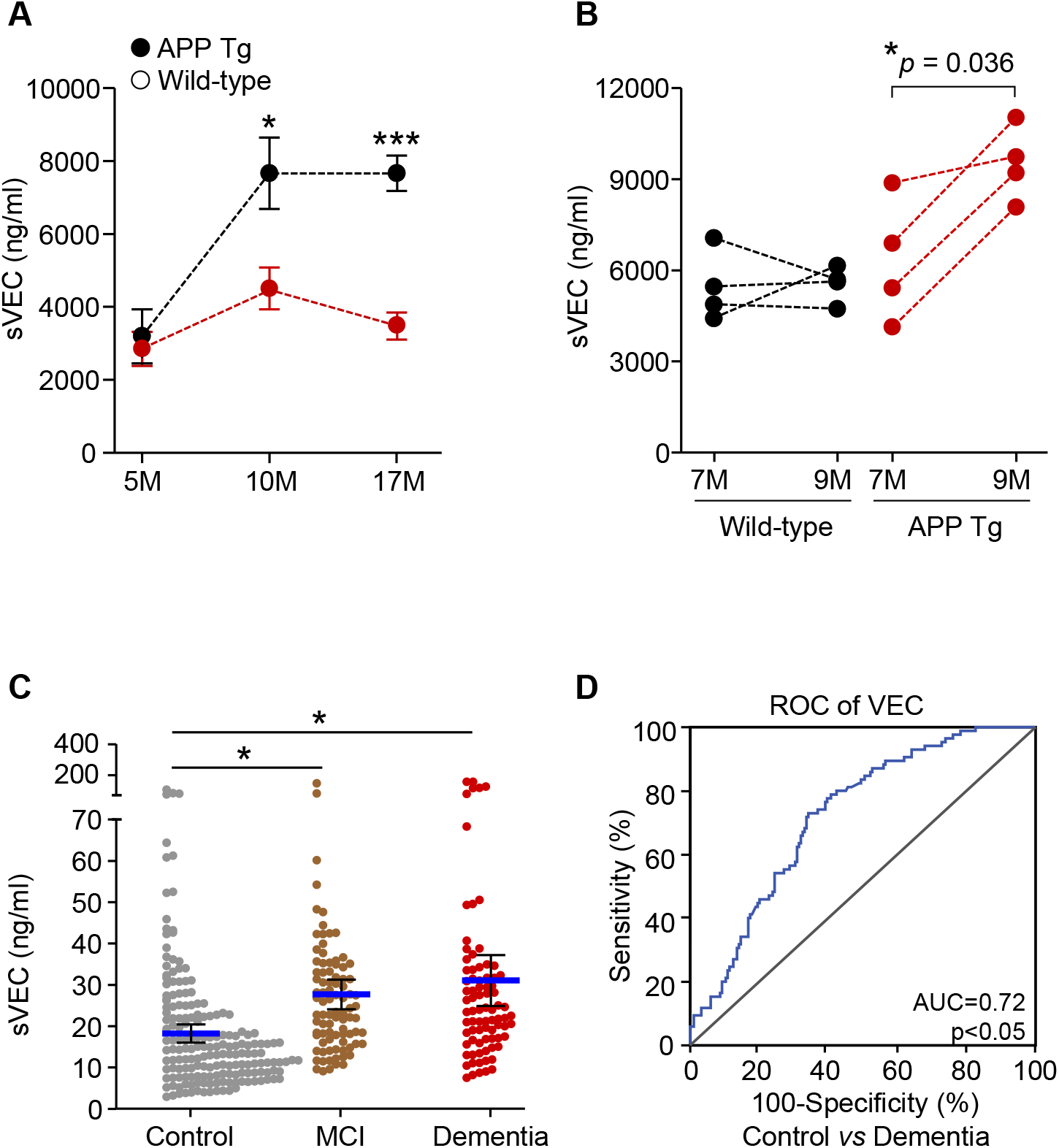
sVEC concentrations in plasma of AD mouse model and human patient. **A**. Cross-sectional analysis of plasma sVEC levels in APP Tg mice and littermates. Plasmas from age-matched mice 5-month-old (n = 3), 10-month-old (n = 7), and 17-month-old (n = 6) were collected. Data were either initially analyzed by ANOVA followed by Turkey posttest. \**P* < 0.05, \*\*\**P* < 0.0001. **B**. Longitudinal analysis of plasma sVEC levels in APP Tg mice and littermates. Column statistic’s with before-after for follow-up to paired data that 7 and 9 months old. Plasma levels as fold change from baseline by 7 month-old wild type. (n = 4). **C**. The levels of plasma sVEC in human patients were measured by ELISA. Mean and 95% confidence interval (CI) of the ratio plasma sVEC for each diagnostic group. Determination of plasma sVEC concentration by ELISA in control (n = 182), MCI (n = 89), and dementia patients (n = 81). \**P* < 0.001, Mann-Whitney U-tests. **D**. Receiver operator characteristic (ROC) curve analysis for the control vs dementia patients. ROC curves obtained from the verification study (set1+set2). The area under the curve and significance levels for the sVEC are indicated

## Discussion

As cerebrovascular lesions in white matter have been observed in AD, microvascular lesions are also common pathological events in AD (Brun & Englund, 1986; White et al, 2002). The main finding of this study was that VEC levels were decreased in the brain capillaries of AD patients and APP Tg mice. The plasma levels of sVEC were increased in both APP Tg mice and subjects with dementia or MCI compared to cognitively normal controls.

BBB hyperpermeability is induced by thrombin, Aβ, and intracellular Ca^2+^ where direct interaction with brain endothelial cells, called BBB opening (Alvarez & Teale, 2006; Brown et al, 2004; Lee et al, 1997; Mackic et al, 1998). As BBB opening is postulated to be dangerous in most circumstances, it may promote AD pathogenesis. Disruption of the BBB is associated with changes in brain endothelial phenotypes and TJ/AJ complex remodeling (Stamatovic et al, 2008). BBB dysregulation has been observed in animal models (Ujiie et al, 2003) and is a clinical feature of AD in human patients (Farrall & Wardlaw, 2009). Although the initiation of BBB opening in AD is unclear, the association of Aβ with BBB disruption has been demonstrated in several mouse models of AD (Ujiie et al, 2003). Therefore, it is possible that amyloidogenesis promotes BBB disruption in AD at the level of the AJ proteins that maintain the BBB. Here, we examined the hypotheses that increased BBB leakage is due to dysfunction of AJ proteins, such as VEC. We showed regulation between the AJ in response to Aβ, potentially altering the integrity of the BBB. A detailed understanding of the mechanisms underlying Aβ-mediated BBB disruption is critical to designing therapeutics directed at reducing neuroinflammation or the development of plaques.

Cadherins are Ca^2+^-dependent transmembrane cell-adhesion molecules, which mediate homophilic interaction between adjacent cells (Angst et al, 2001; Camand et al, 2012). Functional studies of each adhesion molecule might provide a clue on its role during AD development. In neuron, FAD mutations may regulate CREB-mediated transcription by Ncadherin cleavage (Marambaud et al, 2003). It is noteworthy that the brain capillaries obtained from AD brain tissues exhibited significantly lower levels of VEC relative to control human brain tissues. The integrity of the BBB was assessed by investigating AJ morphology in APP Tg mice of various ages; i.e., 2, 5, 9, 12, or 21 months old. 21-month old APP Tg mice were found to have significantly reduced VEC levels, which were correlated with increased BBB opening, while 9-month-old mice tended to show increased leakage of VEC, albeit not significantly so (Fig EV3). These observations suggest that VEC is disappeared in brain vessels and could be an indicator in the early stages of cerebrovascular damage. To our knowledge, this is the first study to provide direct evidence of the disappearance of VEC associated with cerebrovascular damage.

Growing evidence point out the close relationship between cadherins and PS1/γ-secretase in brain. It has been reported that PS1 binds to N-and E-cadherin at the synapse (Baki et al, 2001). PS1/γ-secretase could be also related to disassembly of adherence junction, maintenance, and integrity of brain vasculature. Although a number of studies indicated that TJs are vulnerable to matrix metalloproteinases (MMPs) or oxidative stress (Lum & Roebuck, 2001), little is known about the role of PS1/γ-secretase in the pathological process of neurovascular damage, such as BBB disruption during the development of AD. We have previously reported that PS1/ γ-secretase contributes to AJ disassembly (Jo & Koh, 2013; Park et al, 2008). It was recently reported that PS1/γ-secretase inhibitor reduced the middle cerebral artery occlusion (MCAO)-induced BBB permeability, whereas MMP inhibitors were ineffective (Zhang et al, 2013). Notably, we also showed that PS1/γ-secretase inhibitor blocked Aβ-mediated permeability in EC. The tyrosine phosphorylation of the cytoplasmic domain of VE-cadherin is also involved in the VE-cadherin internalization and increased permeability of EC (Dejana et al, 2008). However, we also rule out the possibility that sVEC production by tyrosine phosphorylation of VE-cadherin because VE-cadherin tyrosine phosphorylation after Aβ treatment was not increased. Given that PS1/γ-secretase is involved in the Aβ-mediated VEC cleavage in endothelial cells, our findings are significant the light of the important role of PS1/γsecretase in alteration of cerebrovascular function. Our observations suggest that PS1/γsecretase may contribute to not only Aβ generation by APP processing but also to cerebrovascular damage.

The results of the present study suggested that sVEC may be useful as a blood marker for clinically diagnosed dementia together with other known markers. In the present, we first showed that blood sVEC is increased in aged APP Tg mice and dementia patients. These observations suggest that the alteration of sVEC levels in blood may be a useful indicator reflecting cerebrovascular damage during the development of AD. Our findings support those of another study in which increased levels of VCAM-1 or soluble PECAM-1 in the blood of AD patients were suggested to reflect dysfunction of the endothelium (Nielsen et al, 2007). In addition, we found that plasma sVEC levels were significantly increased in MCI. The AUC of ROC curve on MCI versus control using plasma sVEC is 0.73 (data not shown). It is interesting that the plasma levels of other endothelial dysfunction markers, VCAM-1 and ICAM-1, were not changed in MCI. Recently, it was reported that soluble VE-cadherin is involved in endothelial barrier breakdown (Flemming et al, 2015). Taken together, Aβ-mediated sVEC production might be involved endothelial barrier breakdown in the early stage of AD progression. Furthermore, we described the presence of sVEC in plasma from dementia patients and investigated its potential clinical relevance in AD. These results suggest that the human plasma VEC concentration could be a useful biomarker of in AD. However, whether the increased sVEC level contributes to the pathogenesis of AD or the high sVEC level is a consequence of disease progression remains to be determined.

Over the last decade, many reports have indicated that fragments of protein in blood may be useful as biomarkers associated with a number of Alzheimer’s-like pathologies. However, there have been few studies regarding blood levels of microvascular markers in AD patients. Although the endothelial dysfunction markers, such as ICAM-1 or VCAM-1 are also found in the circulation and increased levels have been reported in AD (Rentzos et al, 2004; Zuliani et al, 2008), their significance is unclear. Recent DCE-MRI neuroimaging in the lining human brain and CSF findings have demonstrated an early BBB breakdown in patients with mild dementia which correlates pericyte injury as shown by increased levels of pericyte biomarkers (sPDGFRbeta) in the CSF (Sweeney et al, 2015). It should be noted that it is currently unclear which biomarkers reflect BBB dysfunction. As most studies examining microvascular alterations during the development of AD were focused on inflammatory processes, different types of blood-based biomarkers as measures of cerebrovascular damage could be of interest. Elevation of sVEC may indicate VEC cleavage and endothelial dysfunction, which may represent evidence of cerebrovascular damage and BBB opening. Our results supported the possibility of large differences in sVEC as a potential biomarker, allowing clinical validation of its quantification. These observations could aid in better understanding whether brain endothelial function is involved in cognitive decline with vascular risk factors. To our knowledge, this is the first study to suggest that VEC may be a useful microvascular blood biomarker reflecting BBB damage in AD.

In conclusion, our observations suggest that VEC, a protein expressed exclusively in ECs, is subjected to structural modifications in AD (Fig 6). These modifications may be valuable as biomarkers in the AD brain because of the important roles of VEC in vascular permeability as well as BBB disruption. Further studies regarding the relationships between peripheral sVEC and cognitive or other clinical measures of AD will provide further insight into the adequacy of sVEC as a biological marker for this disease.

**Figure 6.**
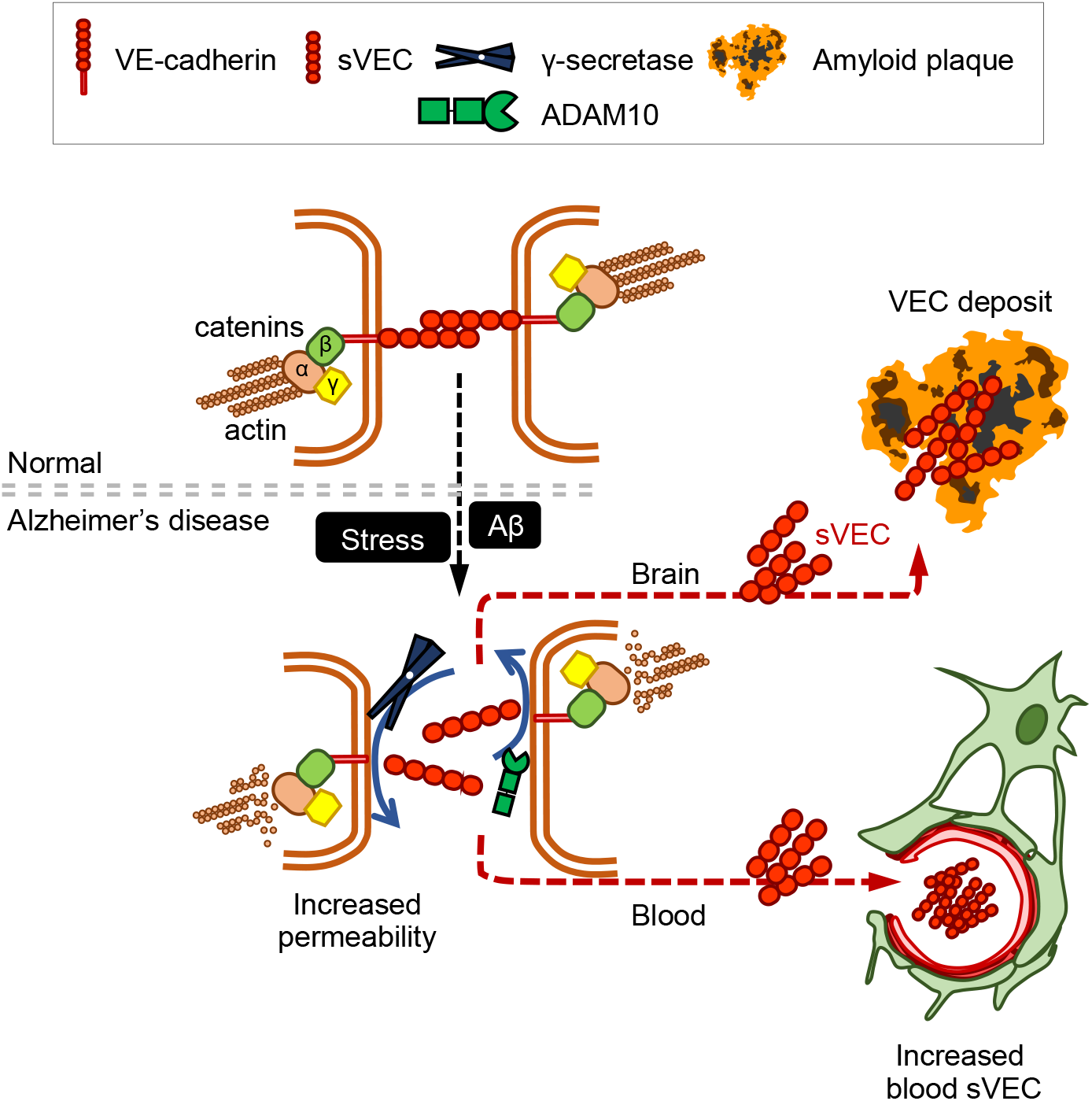
Proposed mechanisms that are likely to be involved in the BBB damage in Alzheimer’s disease. **A**. Amyloid accumulation leads to VEC cleavage and VEC is disappeared at AJ. Soluble VEC (sVEC; VE-cadherin cleavage products) is elevated in blood and deposited with amyloid in brain.

## Materials and Methods

### Subjects

The controls, Mild Cognitive Impairment (MCI) and dementia subjects were selected from the population-based Ansan Geriatric (AGE) cohort established in 2002 to study common geriatric diseases of elderly Koreans aged 60 to 84 years. The sampling protocol and design of the AGE Study have been previously described (Han et al, 2009; Petersen et al, 1999). Cognitive functioning and memory impairments of subjects were assessed using a Korean version of Consortium to Establish a Registry for Alzheimer’s Disease (CERAD-K) neuropsychological battery (Lee et al, 2002). The basic structures of all measures in the original CERAD batteries were maintained in Korean translation. All participants were clinically evaluated according to published guidelines, and each dementia patient met the criteria for the Diagnostic and Statistical Manual of Mental Disorders, fourth edition (Association, 1994). All dementia patients met the criteria for probable AD established by the National Institute of Neurological and Communicative Disorders and Stroke and the Alzheimer’s Disease and Related Disorders Association (NINCDS–ADRDA)(McKhann et al, 1984). MCI was diagnosed on the basis of the Mayo Clinic criteria (Petersen et al, 1999) as described previously (Jang et al, 2010; Kim et al, 2007). The study flow diagram in Fig EV11A provides further details on sample use. In total, blood samples from 352 subjects were included, and the distribution of control, MCI, AD subjects for the Set1 (validation study) and Set2 (extended study) are shown in Table EV1. The study subjects consisted of 81 dementia patients (average age 75.7 ± 5.9, 20 males, 61 females), 89 subjects with MCI (average age 73.7±4.9, 36 males, 53 females), and 182 unrelated healthy controls (average age 70.47±4.9, 86 males, 96 females). Table EV1 summarizes demographic and clinical measures for all covariates tested here, including diagnosis (normal, MCI, dementia), the mini-mental state exam (MMSE), and global clinical dementia rating (CDR). CDR scores are 0 for normal, 0.5 for questionable dementia, 1 for mild dementia, 2 and 3 for moderate to severe dementia (Morris, 1993). The institutional review board of Korea Centers for Disease Control and Prevention (KCDC) approved the research protocol and written informed consent was obtained from all subjects after the nature of the study and its procedures had been explained. Blood samples were collected after informed consent was obtained.

### Human brain tissues

Neuropathological processing of control and AD human brain samples followed the procedures previously established for the Boston University Alzheimer’s Disease Center (BUADC). Institutional Review Board approval for ethical permission was obtained through the BUADC. Next of kin provided informed consent for participation and brain donation. The study was performed in accordance with principles of human subject protection in the Declaration of Helsinki. Detailed information of brain tissues is described in Table EV2.

### Antibodies and reagents

Unless otherwise specified, reagents were obtained from Sigma Aldrich (St Louis, MO, US). The following antibodies were used: mouse monoclonal anti-VE-cadherin extracellular fragment (clone BV9, Abcam), rabbit anti-VE-cadherin cytoplasmic domain (Cell Signal Technology), rabbit anti-phospho VE-cadherin (Y731, GeneTex), rabbit anti-phospho VE-cadherin (Y685, Abcam), rabbit anti-phospho VE-cadherin (Y658, Invitrogen), mouse anti-APP (6E10, Covance), rabbit anti-collagen type IV polyclonal (Fitzgerald), rabbit anti-zonula occludens 1 (ZO1, Invitrogen), rabbit anti-occlaudin (Abcam), rabbit anti-claudin-5 (Abcam), rabbit anti-Aβ_1–42_ (Millipore, AB5078P), anti-presenilin1 (Affinity BioReagents), anti-nicastrin (Affinity BioReagents), anti-Pen-2 (calbiochem). Secondary antibodies for immunofluorescence were donkey antibodies to the appropriate species conjugated with Alexa-Fluor 488, 568 or 647 (1:400, Molecular Probes). The γ-secretase inhibitor DAPT (N-[N-(3,5-difluorophenacetyl)-L-alanyl]-(S)-phenylglycine t-butyl esterand), 1,2-bis(o-Aminophenoxy)ethane-N,N,N′,N′-tetraacetic acid tetra (acetoxy- methyl)ester (BAPTA-AM), MMP9 inhibitor GM6001 and 4′,6-diamidino-2-phenylindole (DAPI) were purchased from Calbiochem. The ADAM10 inhibitor GI254023X was purchased from Sigma-Aldrich. Aβ_1–40_ (Invitrogen), Aβ_1–42_(Invitrogen), and nonamyloidogenic scrambled Aβ_42–1_ (Biosource) peptides were dissolved in hexafluoroisopropanol for 2 h at room temperature, and lyophilized peptide was dissolved in dimethylsulfoxide (DMSO).

### Animals

APP transgenic mice were used in the present study, as previously reported (Yun et al, 2013). All studies were conducted with a protocol approved by the local Institutional Animal Care Use Committee in compliance with Korea National Institute of Health guidelines for the care and use of experimental animals.

### ELISA measurements

All the cell-free plasma samples were stored in aliquots at –80°Cuntil assayed collectively by an investigator who was blinded to patient assignment. ELISA was carried out with the human VE-cadherin and VCAM-1 ELISA kit (Bender Medsystems GmbH, Vienna, Austria) and ICAM-1 ELISA kit (R&D System, Heidelberg, Germany) according to the manufacturer’s instructions. Calibrations were carried out duplicates, intra- and inter-assay CVs within the range given by the manufacturers, respectively. All experiments were performed by an investigator blinded to the study groups assignment.

### Immunoperoxidase staining

Brain from APP Swedish/PSEN1dE9 mice together with their littermate controls were fixed in 4 % (wt/vol) paraformaldehyde. Cryostat sagittal sections were cut on a sliding into 16 μm slices at –80°C and placed on a microslide for immunostaining. The sections were immunostained with 6E10 (1:1000; Covance), n-terminus VE-cadherin antibody (1:100; BV9, Abcam), c-terminus VE-cadherin antibody (1:200; a.a.770–781, ECM Biosciences), PECAM-1 (1:250; BD Bioscience) vWF (1:400, Origene), and Collagen-iv (1:200; Fitzgerald). Sections were then incubated for 1 h at RT with secondary antibodies conjugated biotinylated. After then Avidin/Biotin complex reagents use increased sensitivity. 3’3-diaminobenzidine in chromogen solution to visualize. VE-cadherin /vWF stained sections were counterstained with thioflavin-S, clear and mount. Axiolab-Pol polarizing (Carl Zeiss) microscopy with Axio Vision Release 4.8 software was used for analysis of 3,3’-Diaminobenzidine (DAB) photomicrograph and localization of proteins. Images were analyzed with the NIH Image J. Imaging for immunofluorescence was performed on a fluorescence microscope and image capture.

### Immunofluorescence

For labeling, 18 μm mounted section was blocked for 1 h in 5 % BSA, 0.03 % Triton X-100 in PBS and incubated overnight at 4°Cwith primary antibodies in 3 % BSA, 0.03 % Triton X-100 in PBS. Sections were washed with PBS and developed with an anti-species specific Alexa Fluorconjugated secondary antibodies (Moecular probe). Sections were washed with PBS. Tissue section slides were mounted with or without DAPI (Vector Laboratories). Fluorescent photomicrographs were captured using Zeiss microscope or confocal microscope and exported to Ziess LSM. Image was analyzed with the NIH Scion image.

### Statistical analyses

Results are expressed as mean ± SEM. Significance was analyzed using two-tailed Student’s *t-* test and ANOVA. To compare demographic and plasma biomarkers data between dementia, MCI, and control groups, Kruskal-Wallis test was performed followed by Mann-Whitney U-tests. Correlation between factors was analyzed by Spearman’s method. A chi-square test was used to compare the sex variation of AD, MCI, and control groups. Statistical analyses were conducted using SPSS for Windows (SPSS Inc.) or R (Bell Laboratories, version 3.1.1). Values of *P* < 0.05 were considered statistically significant.

## Acknowledgments

We thank Mr. Jae Chun Song for maintaining the facilities and equipment used in the animal experiments. This research was supported by a fund (2012-NG62003–00, 2016-NG62002–00, 2017-NI62001–00) by Research of Korea Centers for Disease Control and Prevention.

## Author contribution

D.H.L. and Y.H.K. designed the study, analyzed the data and wrote the manuscript. S.J.C., H.J.L., and C.J. performed the cell biology experiments. J.W.S performed the statistical analyses. S.A.J., M.H.P. and C.H. designed the study, provided clinical samples and analyzed the data. N.K. and H.R contributed to study design and analyzed the data for human brain tissues. R.E.T. contributed to study design and discussion. All authors contributed to analysis and discussion of the results.

## The pater explained

### Problem

Cerebrovascular abnormalities may contribute to the pathology of Alzheimer’s disease (AD). *In vivo* assessment of microvascular pathology provides a promising approach to develop useful biological markers for early detection and pathological characterization of AD. Although impairment of blood brain barrier by Aβ deposits leads to disintegration of endothelial junctional and an increase in permeability of the blood–brain barrier (BBB), little is known about the molecular mechanisms regulating the cohesion of endothelial adherens junctions (AJs) in AD.

### Results

We have investigated BBB disruption and the role of VE-cadherin in the progression of amyloid pathology. We show that initial increase in cerebral Aβ deposits leads to microvascular pathology provide alteration of VEC integrity in BBB *in vitro* and *in vivo*. Our data provide evidence for alteration of VEC in the brain of AD patients. In addition, soluble VE-cadherin (sVEC) levels in blood were elevated in and Mild Cognitive Impairment (MCI) and dementia patients compared with control.

### Impact

Our finding suggests that alteration of VE-cadherin may reflect the attenuation of BBB integrity in AD. Plasma VE-cadherin may have the potential to detect AD earlier than microvascular pathology in AD or Aβ burden test.

